# Heavy Metal Exposure Disrupts Electrotactic Behavior in *Caenorhabditis elegans*

**DOI:** 10.1101/2025.05.20.655158

**Authors:** Shane K. B. Taylor, Muhammad H. Minhas, Justin Tong, P. Ravi Selvaganapathy, Ram K. Mishra, Bhagwati P. Gupta

## Abstract

Environmental toxicants such as heavy metals can profoundly impact organismal physiology, yet sensitive and rapid behavioral assays to quantify such effects are limited. Here, we employed a microfluidic-based electrotaxis assay to systematically evaluate the impact of chronic exposure to metal salts on the electrotactic swimming behavior of the nematode *Caenorhabditis elegans*. We report that exposure to Ag, Hg, MeHg, Cu, Mn, and Pb significantly alters electrotaxis speed. Notably, Cu required higher concentrations to induce phenotypes, whereas Ag and Hg disrupted behavior at lower doses. Three other metals (Ni, Fe, and Cd) did not elicit marked electrotaxis defects. Bioaccumulation analysis of Cu, Ag, Hg, and MeHg via ICP-MS revealed that MeHg was present in highest amount in worms. Consistent with this, MeHg showed the highest toxicity. We also found dopaminergic neurodegeneration in most metal-exposed animals, correlating with behavioral impairments. Although heat shock factor-1 (HSF-1) and its downstream target *hsp-16.2* were induced by some metals, their expression did not consistently correlate with electrotaxis defects. Moreover, metal-induced impairments persisted even after recovery on toxin-free media, indicating possible irreversible damage. Our findings establish electrotaxis in *C. elegans* as a robust, non-invasive assay to assess neurobehavioral toxicity and demonstrate its utility for detecting sub-lethal impacts of environmental metals.

## INTRODUCTION

Environmental exposure to heavy metals poses a significant threat to human and ecological health. These toxicants can originate from diverse sources including industrial discharge, contaminated food and water, and airborne particulates, making them pervasive in daily life. Organisms have evolved multiple detoxification strategies such as restricting metal uptake, enhancing efflux, or activating stress-responsive pathways to mitigate their harmful effects. While mammalian models such as mice and rats offer valuable insights into metal toxicity, their use is limited by complexity, cost of maintaining cultures, and ethical considerations. Similarly, in vitro systems like cell cultures fail to replicate organismal-level responses. These limitations have driven the adoption of alternative model systems such as the nematode (worm) *Caenorhabditis elegans.* As an invertebrate system, *C. elegans* offers numerous advantages including genetic tractability, rapid lifecycle, low maintenance costs, and a simple, well-characterized nervous system.

*C. elegans* are sensitive to most toxicants of interest, including metal salts, and have been used for toxicity studies (Hunt, 2017). Its responses are often observed at environmentally relevant concentrations, making it a useful bioindicator for toxicity screening (Akinyemi et al., 2019; Ju et al., 2013; Wu et al., 2012). In this study, we report the impact of chronic exposure to eight heavy metals, mercury (Hg), silver (Ag), copper (Cu), iron (Fe), lead (Pb), manganese (Mn), nickel (Ni), and cadmium (Cd), on the electrotactic behavior of *C. elegans*. Electrotaxis, or directed movement in response to a direct current (DC) electric field, offers a robust behavioral readout with minimal variability, ideal for detecting subtle neuromuscular impairments (Rezai et al., 2010).

Among these metals, Hg and its organometallic derivative methylmercury (MeHg) are especially notorious for their neurotoxicity. MeHg accumulates in aquatic systems through methylation processes and poses a grave risk due to its ability to cross biological membranes and target neural tissues (Forsyth et al., 2004; Hong et al., 2012). Hg is most infamous for its neurotoxic properties: even acute exposures can induce an assortment of cognitive, personality, sensory, and motor abnormalities, possibly culminating in coma and death (Clarkson and Magos, 2006). Other metals such as Cu, Fe, and Mn are essential micronutrients involved in critical enzymatic reactions. However, their excess is harmful, leading to oxidative stress, mitochondrial dysfunction, and neurodegeneration (Kumar et al., 2015b; Ndayisaba et al., 2019; Raj et al., 2021; Wazir and Ghobrial, 2017). Experiments in mice have found that bioaccumulation of Mn affects GST and AChE activity, lipids and proteins. Similar to Cu and Mn, Fe also plays important roles in healthy organisms; yet, acute Fe overload can cause iron poisoning (Chen et al., 2013).

Ag, Ni, Pb, and Cd, although non-essential, are widely present in industrial and consumer products. Ag, for example, can cause dermal and ocular toxicity, and its prolonged exposure leads to argyria and argyrosis (Drake and Hazelwood, 2005; Hadrup et al., 2018). Other studies on Ag toxicity have reported developmental deformities in zebrafish, neurotoxicity in mice, and mitochondrial damage to human cells (Asharani et al., 2009; Bar-Ilan et al., 2009; Gonzalez-Carter et al., 2017; Rahman et al., 2009). Similarly, Ni, which is fairly ubiquitous in industrial and consumer products, is associated with toxicity at large doses or with chronic exposure. Chronic exposures to Ni affect skin such as causing itching or rashes, respiratory defect, and some forms of cancers affecting lung and other tissues (Genchi et al., 2020). Experiments in mice have shown that Ni causes acute lung inflammation (Mo et al., 2019), although skin and cancerous phenotypes were not observed. Pb is a pervasive environmental neurotoxicant; it is particularly toxic to developmental brains, causing long-term detrimental effects on learning, memory and behavior in children (Neal and Guilarte, 2010). Pb poisoning also results in symptoms such as neuromuscular weakness., anemia, delirium, convulsions, and death (Jaishankar et al., 2014). Studies have shown that high Pb exposures in mice can lead to altered movement, exploratory responses, and hyperactivity (Jaishankar et al., 2014; Silbergeld and Goldberg, 1974). And, finally, Cd is a known carcinogen that has been associated with mutagenic responses (Jaishankar et al., 2014). Cellular and molecular studies have report that Cd exposures affect expression of genes involved in cell cycle, apoptosis, oxidative stress and inflammation (Kim et al., 2017). Studies on Cd exposures have also reported cirrhosis, kidney dysfunction, and defects in the skeletal system (Jaishankar et al., 2014).

Standard behavioral assays in *C. elegans*, including measurements of locomotion, viability, fertility, and lifespan, have long served as effective endpoints in toxicology (Chen et al., 2013; Tang et al., 2019; Wang and Xing, 2008). Moreover, the use of transgenic lines and fluorescent microscopy enables visualization of metal-induced damage (Chen et al., 2013). Locomotory behavior is a well-established endpoint that serves as a simple yet highly reliable indicator of toxicity, with slow and abnormally uncoordinated movement being associated with nervous system defects (Hart, 2006). However, traditional assays are often hindered by spontaneous behavioral variability such as pausing, reversals, and reorientations, which complicates data interpretation.

To address these limitations, we employed a novel microfluidic-based electrotaxis assay, which enables precise, reproducible quantification of movement behavior under a constant directional stimulus. We previously showed that this assay provides a sensitive and high-throughput method for analyzing behavioral phenotypes in *C. elegans* (Rezai et al., 2010; Tong et al., 2013). Here, we use this platform to systematically investigate how chronic exposure to eight heavy metals alters electrotactic behavior. We find that Ag, Hg, MeHg, Cu, Mn, and Pb impair electrotaxis in distinct ways, while Ni, Fe, and Cd do not. Further analysis reveals a correlation between behavioral defects and dopaminergic neurodegeneration for most metals tested, which is consistent with previous findings of dopamine signaling being involved in mediating electrotaxis behavior (Salam et al., 2013). We carried out additional experiments to investigate the basis of electrotaxis defects. The analysis of cytosolic unfolded protein response (UPR) did not show a correlation between metal toxicity, stress response, and electrotaxis, suggesting differences in how metals affect behavior. Notably, metal-induced defects often persisted after the cessation of exposure, indicating lasting or possibly irreversible damage.

Our study represents the first application of electrotaxis as a behavioral endpoint for assessing chronic metal toxicity. The findings not only highlight electrotaxis as a rapid and quantitative assay but also provide insights into the differential toxic effects of various heavy metals on neuromuscular function in *C. elegans*.

## RESULTS

### Chronic exposures to some metal salts cause defects in electrotaxis

To evaluate the toxic effects of metal exposure on the electrotaxis behavior of worms, we introduced one-day old adult worms, grown in the presence of metal toxicants, into the microchannel device and measured their electrotaxis speed. Initially, worms were exposed to three different metals, Ag, Hg, and Cu. Previous studies have reported LC50 values for these chemicals (Du and Wang, 2009; Ruszkiewicz et al., 2018; Shen et al., 2009). Based on this, we chose 5 μM as a starting concentration for electrotaxis assays. While Ag and Hg caused reduced electrotaxis speed, Cu had no such effect (**Figure 1A**). Two additional doses, 50 μM and 150 μM, were also tested. For Ag and Hg, both doses were highly toxic thereby precluding electrotaxis assays, whereas Cu once again showed no obvious adverse effect (**Figure 1A**, see Methods). It took a much higher level of Cu, i.e., 500 μM, to elicit an electrotaxis defect (**Figure 1A**). Consistent with these results, examination of other biological endpoints, namely brood size, body length, and lifespan also supported detrimental effects of metals on animals (**Supplementary Figure 1**), and are in line with previous findings (Roh et al., 2009). Specifically, Ag and Cu inhibited reproduction, however Hg caused no visible change. Body length was not affected by Ag and Hg, but Cu exposure led to a small but statistically significant reduction. All metals had harmful effects on the lifespan of animals (**Supplementary Figure 1**).

**Figure 1.**
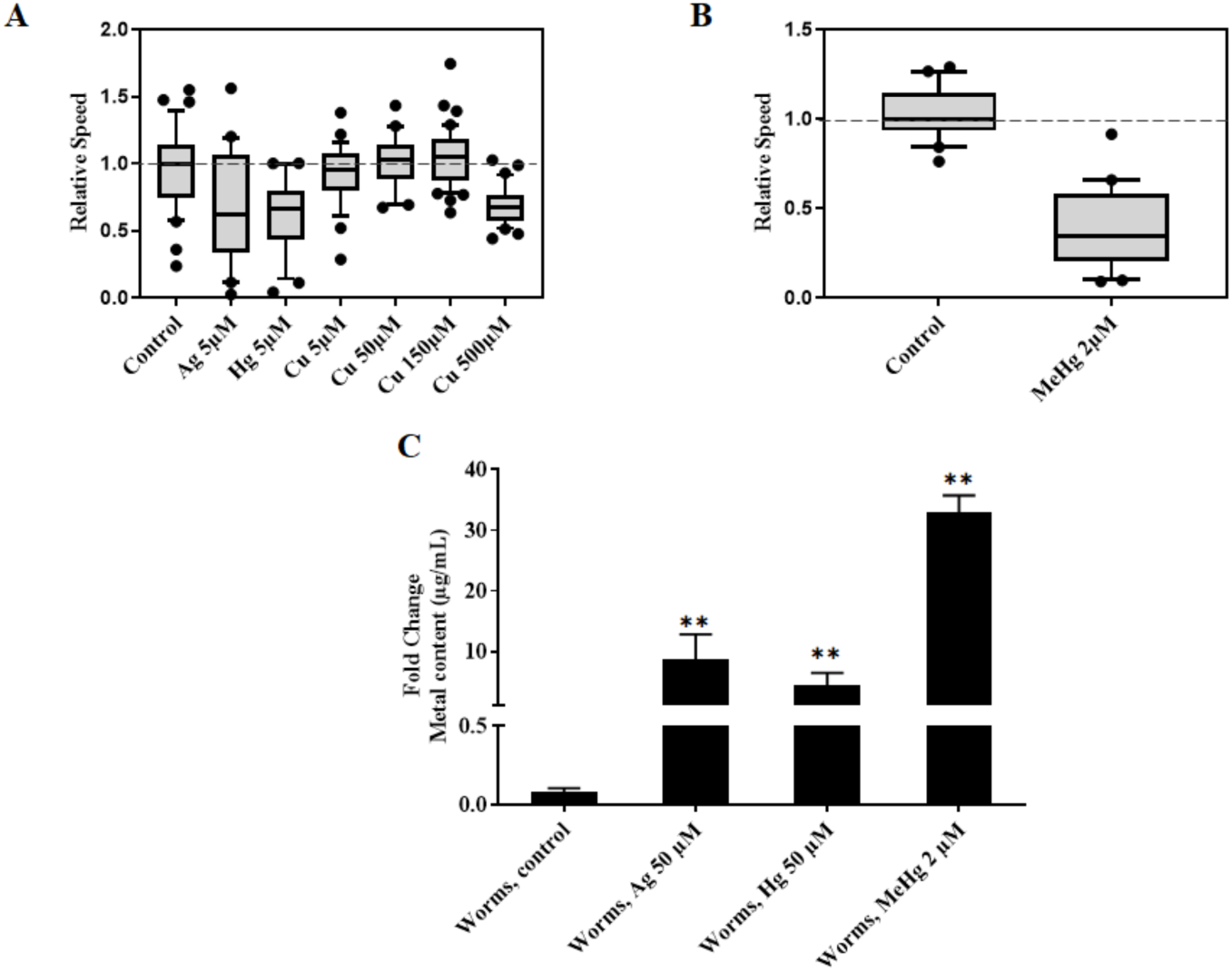
Electrotaxis of animals chronically exposed to Ag, Hg or Cu. Boxes represent measurements from 25^th^ to 75^th^ percentiles, central horizontal lines represent medians, vertical lines extend to 10^th^ and 90^th^ percentiles, and dots represent outliers. **A)** Defects manifesting as slower swimming speeds appear in animals grown on plates containing 5 μM Ag (P < 0.01) or 5 μM Hg (P < 0.01). Cu-induced electrotaxis speed defects do not appear with plates containing 5 μM (P > 0.05), 50 μM (P > 0.05) or 150 μM (P > 0.05) but do appear at 500 μM (P < 0.01). **B)** Defects manifesting as slower swimming speeds appear in animals grown on plates containing 2 μM MeHg (P < 0.01). **C)** Metal content in *C. elegans* following 69 h metal salt exposure. Elemental content was measured as a function of total sample volume. Metal content in worms is significantly higher than in control following treatment with Ag 50 μM ∼5.4 μg/mL, Hg 50 μM ∼10.0 μg/mL, or MeHg 2 μM ∼0.4 μg Hg/mL. Data is pooled from independent replicates (n > 30 animals). Statistical analysis for panel **A** and **C** was done using a one-way ANOVA with Dunnett’s post hoc test. Student’s unpaired t-test was used for panel **B**. Significant data points are marked by the following: *p<0.05; **p<0.01; ***p<0.001; ****p<0.0001. **C** is plotted as mean ± SEM

An organic Hg salt was also tested. As mentioned above, the methyl form of Hg (MeHg) is particularly toxic to humans and causes nervous system abnormalities and birth defects (Hong et al., 2012). *C. elegans* are highly sensitive to MeHg since even a short acute exposure leads to lethality and defects in development, reproduction and locomotion (Helmcke and Aschner, 2010; McElwee and Freedman, 2011). Of the four different concentrations tested (2 μM, 4.5 μM, 9 μM, 18 μM), the lowest dose (2 μM) was the only one suitable for electrotaxis assays as it did not appear to harm the animals at the plate level although progeny counts were significantly reduced (data not shown). Exposed animals showed a major reduction in electrotaxis (**Figure 1B**). Overall, the results show that the electrotaxis phenotype of *C. elegans* is highly sensitive to low, chronic doses of Ag, Hg, and MeHg but a higher dose of Cu. Additionally, movement defects correlate well with other phenotypes observed.

To determine if electrotaxis abnormalities were due to metals being taken up by animals in our exposure paradigm, we used the ICP-MS technique. The analysis revealed that metal contents were significantly higher inside worms than what was recovered from agar media. While Ag, Hg, and Cu accumulated at 8-fold, 4-fold, and 4-fold, respectively, MeHg accumulation was 33-fold (**Figure 1C, Supplemental Figure 2**). These data show that metal toxicants, particularly MeHg, increase at high levels inside animals’ bodies when present in their external environment. The results demonstrate an efficient uptake of metals in our method.

Having demonstrated the effect of three metals (Ag, Hg, and Cu) on electrotaxis, we tested an additional set of five metals, Cd-50μM, Fe-150μM, Ni-150μM, Mn-150μM and Pb-150μM. The concentrations of all of these used in our experiments were well below published LC50 (Du and Wang, 2009; Ruszkiewicz et al., 2018; Shen et al., 2009). Plate-level observations showed that animals were healthy and exhibited responses similar to wild-type animals (see Methods and **Supplementary table 1**). Analysis of the electrotaxis responses showed that Mn and Pb were the only two metals that caused defects (**Figure 2A**). Interestingly, while Mn reduced the speed, Pb exposure resulted in a hyperactive response (**Figure 2A**). The remaining metals (Ni, Fe and Cd) did not cause any change in electrotaxis response of exposed animals.

**Figure 2.**
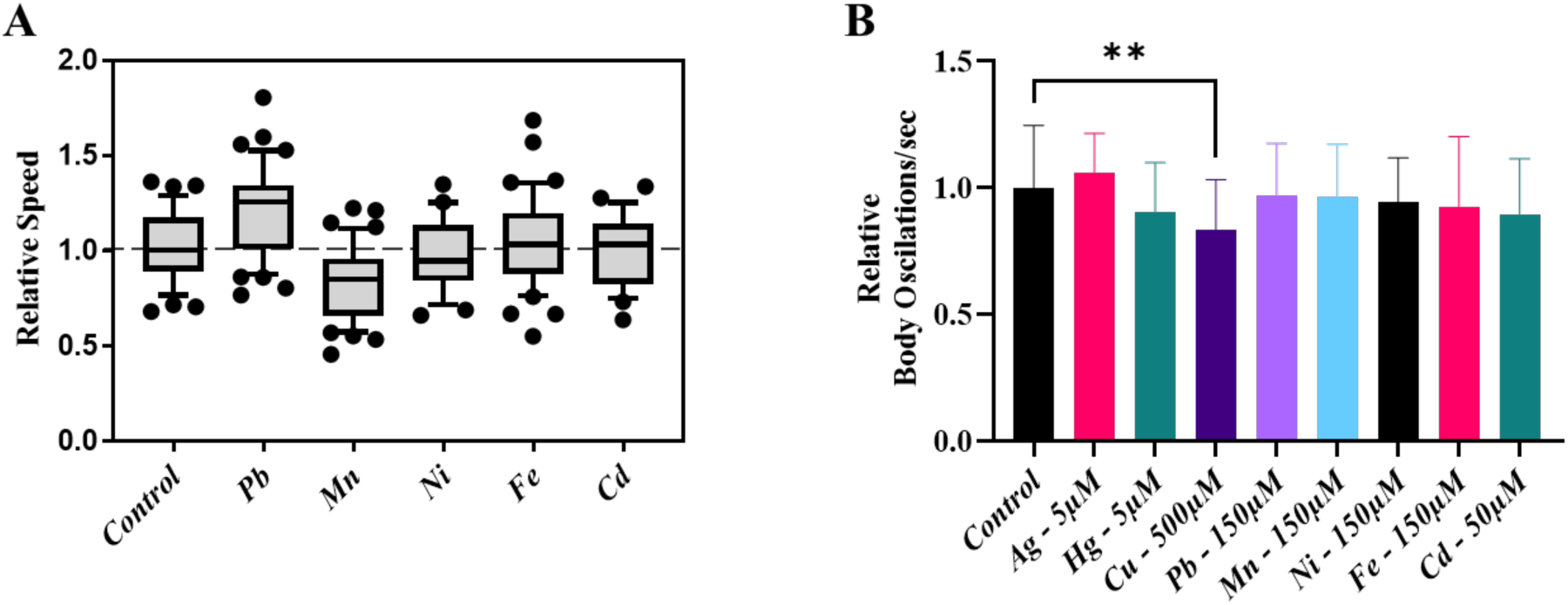
Analysis of movement defects in animals exposed to a set of heavy metals. (**A**) Box plot showing electrotactic responses of animals chronically exposed to Pb, Mn, Cd, Ni, Fe and Cd. Both Cu and Mn caused a slower speed (p <0.001), Pb results in an increased speed (p<0.001). The other metals Ni, Fe, and Cd did not alter the speed of the animals. Refer to Figure 1 for a description of the plot. (**B**) The body oscillations of chronically exposed animals in the microfluidic channel calculated using the MATLab software. Cu had significantly more body oscillations within the channel compared to control. The other metals did affect the body oscillations. Data is pooled from independent replicates of 3 (n=15 to 45 animals). Statistical analysis for panel A and B was done using a one-way ANOVA with Dunnett’s post hoc test. *p<0.05; **p<0.01; ***p<0.001; ****p<0.0001. **B** is plotted as mean ± SD

We also analyzed body oscillations, which is another parameter of movement (Hart, 2006; Wang and Xing, 2008), to see if these correlated with speed defects. The data showed that except for 500 μM Cu, which resulted in slower oscillations, the responses of all metals are comparable to wild type controls (**Figure 2B, Supplementary Figure 3**). A lack of correlation with electrotaxis speed indicates that changes in the electrotaxis speed may not be affected by body bending characteristics of animals, although minor effects cannot be ruled out. Overall, our findings show that the electrotaxis response of animals is sensitive to chronic exposures of Ag, Cu, Hg, MeHg, Mn and Pb.

### Prolonged exposure to metals can cause electrotaxis defect

We reasoned that a lack of phenotype for some metals may be due to insufficient exposure and a longer exposure duration may result in an abnormal response. To investigate this further, we grew worms on culture plates containing Ni (150μM), Fe (150μM) and Cd (50μM) till day 3 of adulthood. In addition, 150μM Cu was tested which had no effect on electrotaxis in our previous assay (see above). The electrotaxis response of animals was examined and the results showed that while Ni, Fe, and Cd had no detrimental effect, Cu exposure caused a significant reduction in speed (**Figure 3**).

**Figure 3:**
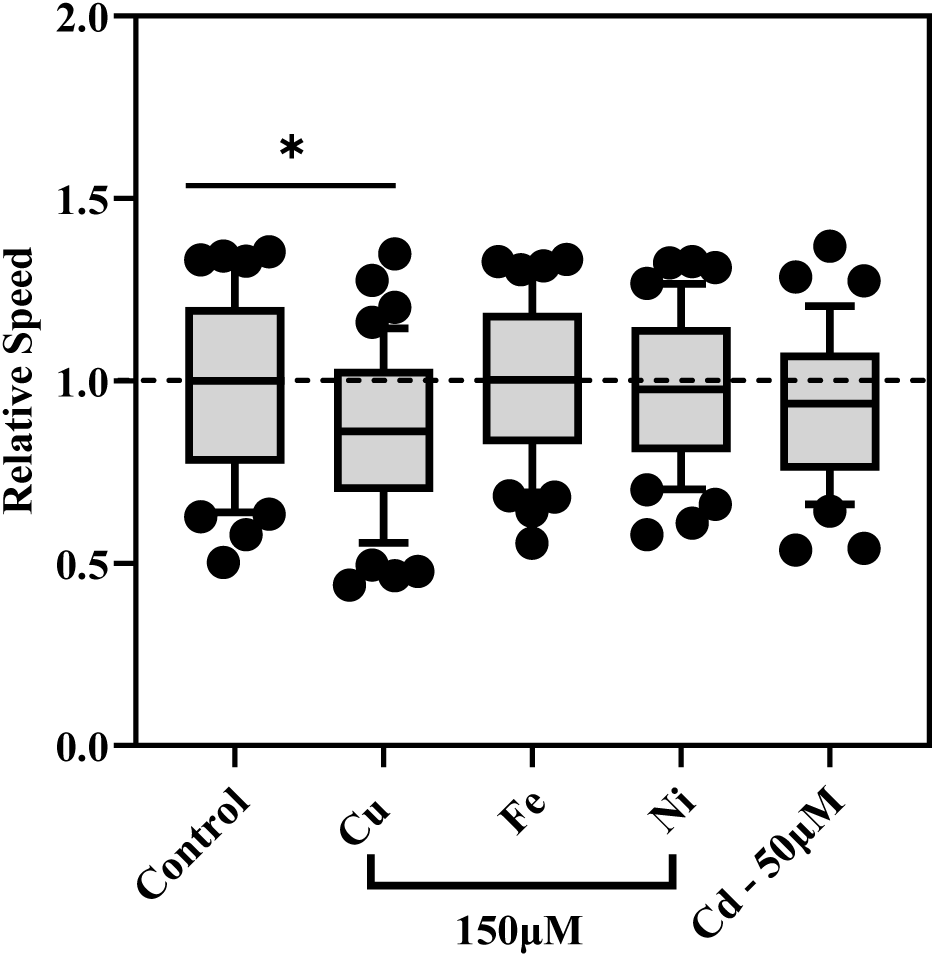
Electrotaxis responses of animals exposed to metal salts. Animals were exposed to four different metal salts until day 3 of adulthood (150 μM Cu, 150 μM Fe, 150 μM Ni, and 50 μM Cd). Only Cu-exposed animals exhibited a significantly reduced speed (p= 0.0275). The other heavy metals Fe, Ni and Cd did not alter the electrotaxis speed of the animals (p= 0.9902,0.9999,0.6746 respectively). Data is pooled from independent replicates (n > 30 animals). Statistical analysis for panel A and B was done using a one-way ANOVA with Dunnett’s post hoc test. *p<0.05; **p<0.01; ***p<0.001; ****p<0.0001.

We also evaluated general growth and movement characteristics of animals on plates and found that while Cd exposure had a mild effect on larval growth, the adults were visually normal. The other three metals (Cu, Ni, and Fe) caused no obvious abnormality and animals appeared healthy and exhibited normal movement and growth characteristics. We conclude that the electrotaxis assay provides a quantitative and sensitive means to examine the detrimental effect of prolonged exposure of Cu on *C. elegans*.

### Effects of metal exposures on DAergic neurons and cytosolic stress response

Since dopamine signaling affects electrotaxis behavior of animals (Salam et al., 2013), we reasoned that metals may affect neuronal phenotypes. This is consistent with other studies showing that heavy metals cause damage to dopaminergic neurons and affect dopamine signaling (Benedetto et al., 2010; Jaishankar et al., 2014). To this end, DAergic neurons were examined using a *dat-1p::YFP* marker. In wild-type animals, the DA transporter *dat-1* is expressed in four pairs of CEP and one pair of ADE neurons, which allows easy identification of cell bodies and their projections (**Figure 4A**). Upon quantification, we found that Ag and Hg caused damage in more than half of the worms examined, Cu affected roughly 40% of the population (**Figure 4A & B**). The phenotypes included loss of cell bodies, and faint and punctate projections. Of the remaining metals, Mn, Pb, and Cd affected roughly one-third of animals, with phenotypes mainly confined to neuronal projections. Ni and Fe did not exhibit any significant neuronal defects. We conclude that most metals cause neuronal damage.

**Figure 4.**
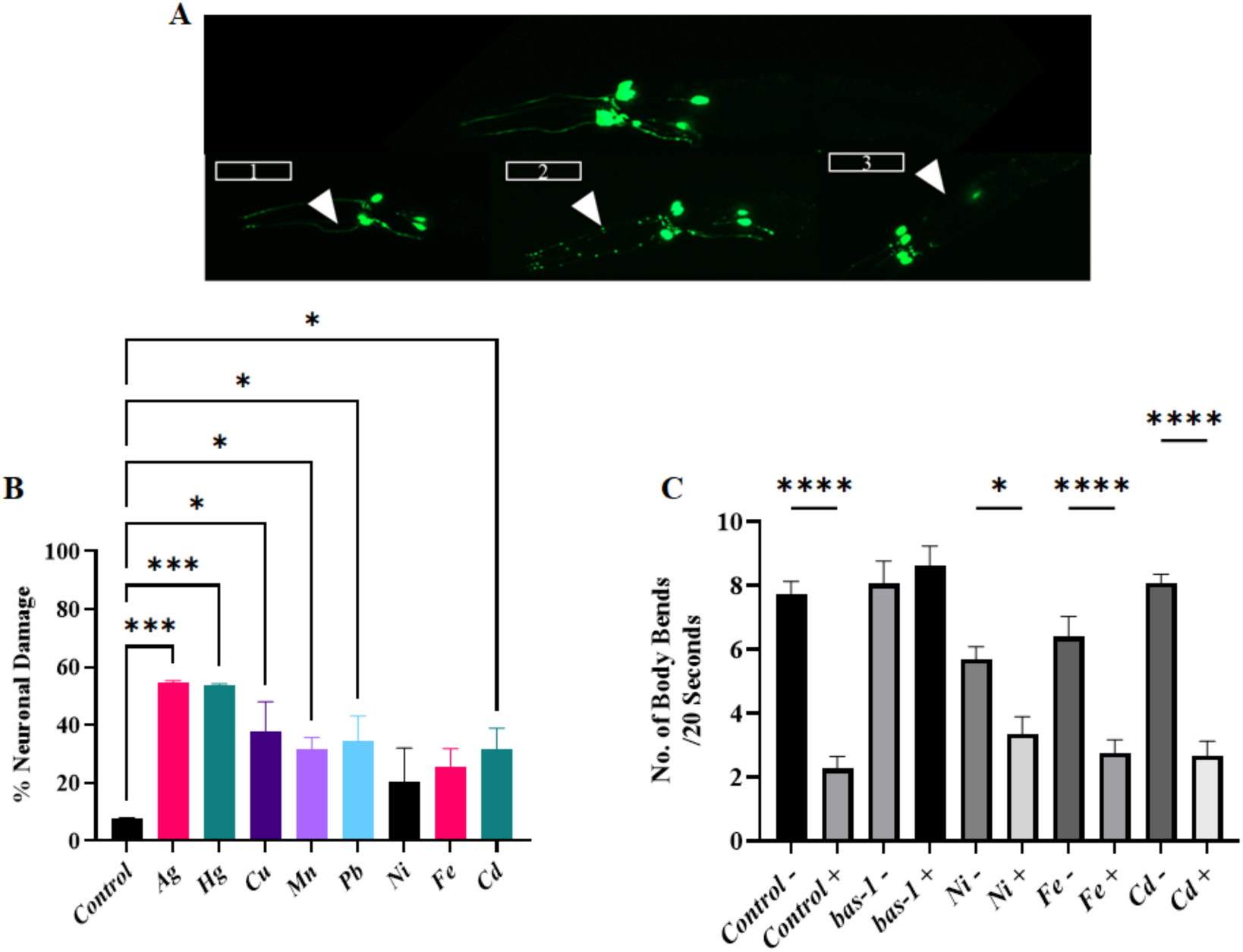
Neurodegenerative phenotypes of young adult *dat-1p::YFP* transgenic animals resulting from chronic exposure to metal salts. **A)** Representative images of neuronal defects observed in animals. Dopaminergic neurons and processes are healthy in untreated *dat-1p::YFP* control animals. Sub-panels show the defects that arise in metal-exposed dat-1p::YFP animals including 1. abnormal CEP dendrite morphology, 2. punctate patterns in CEP dendrites and ectopic neurite growth, and 3. loss of CEP neurons and dendrite segments. **B)** The percentage distribution of *dat-1p::YFP* animals with normal fluorescent neurons following metal exposure. **C)** The Basal slowing response of Day 3 adults following metal Exposure. Well-fed animals exposed to metals were transferred to plates (+) with and (-) without bacteria. As a control both wildtype *N2* and *bas-1* animals were used. *N2* had a normal BSR and as expected *bas-1* animals had a defective BSR, demonstrating the effectiveness of this assay in our hands. Data is expressed as ± SEM and pooled from individual replicates (n=16 animals) per group. Statistical analysis for panel **B** and **C** was done using a one-way ANOVA with Dunnett’s post hoc test. *p<0.05; **p<0.01; ***p<0.001; ****p<0.0001.

Since Ni, Fe, and Cd did not alter electrotaxis and two of these (Ni and Fe) also had no effect on DAergic neurons, we used another behavioral paradigm to examine the animals. The locomotory response of animals was measured in the presence of food using an established basal slowing response (BSR) assay, which is mediated by DAergic neuron signaling (Sawin et al., 2000). Animals having abnormal DAergic neuronal function, such as *bas-1* (DOPA decarboxylase) mutant, have defective BSR (Sawin et al., 2000) (**Figure 4C**). We found that none of the three metals affected the BSR of animals (**Figure 4C**), a finding that is consistent with their electrotaxis phenotypes. Thus, while many of the metals affect electrotaxis and DAergic neurons, both phenotypes do not always co-exist.

Previous work from our lab has shown that chemicals which alter electrotaxis behavior can affect the stress responses of animals (Taylor et al., 2021). Additionally, toxic metals are known to compromise unfolded protein response (UPR) pathways such as the cytosolic heat shock response (HSR) which is known to work with metallothionein in clearing metals leading to reduced cellular stress and proteotoxicity (Liu and Thiele, 1996; Martinez-Finley and Aschner, 2011; Uenishi et al., 2006). To investigate the role of HSR in metal-induced response, we first examined the effect of Mn and Pb, which affect electrotaxis but in an opposite way, and Cd that causes no electrotaxis phenotype but activates the HSR marker *hsp-16.2* (Wang et al., 2020). The qPCR analysis showed that while Mn and Cd increased *hsp-16.2* transcript levels, Pb had no significant effect (**Figure 5A**). Interestingly, there was no increase in *hsf-1* levels, a transcription factor that regulates expression of small heat shock chaperons including *hsp-16.2*. However, Mn and Cd had a slight significant decrease in *hsf-1* expression. It is conceivable that *hsf-1* affects *hsp-16.2* levels by acting post-transcriptionally or additional heat shock proteins may be involved. Moreover, electrotaxis defects in *hsf-1* mutants (Taylor et al., 2021) were not further enhanced by Mn, Pb and Cd (**Figure 5B**).

**Figure 5:**
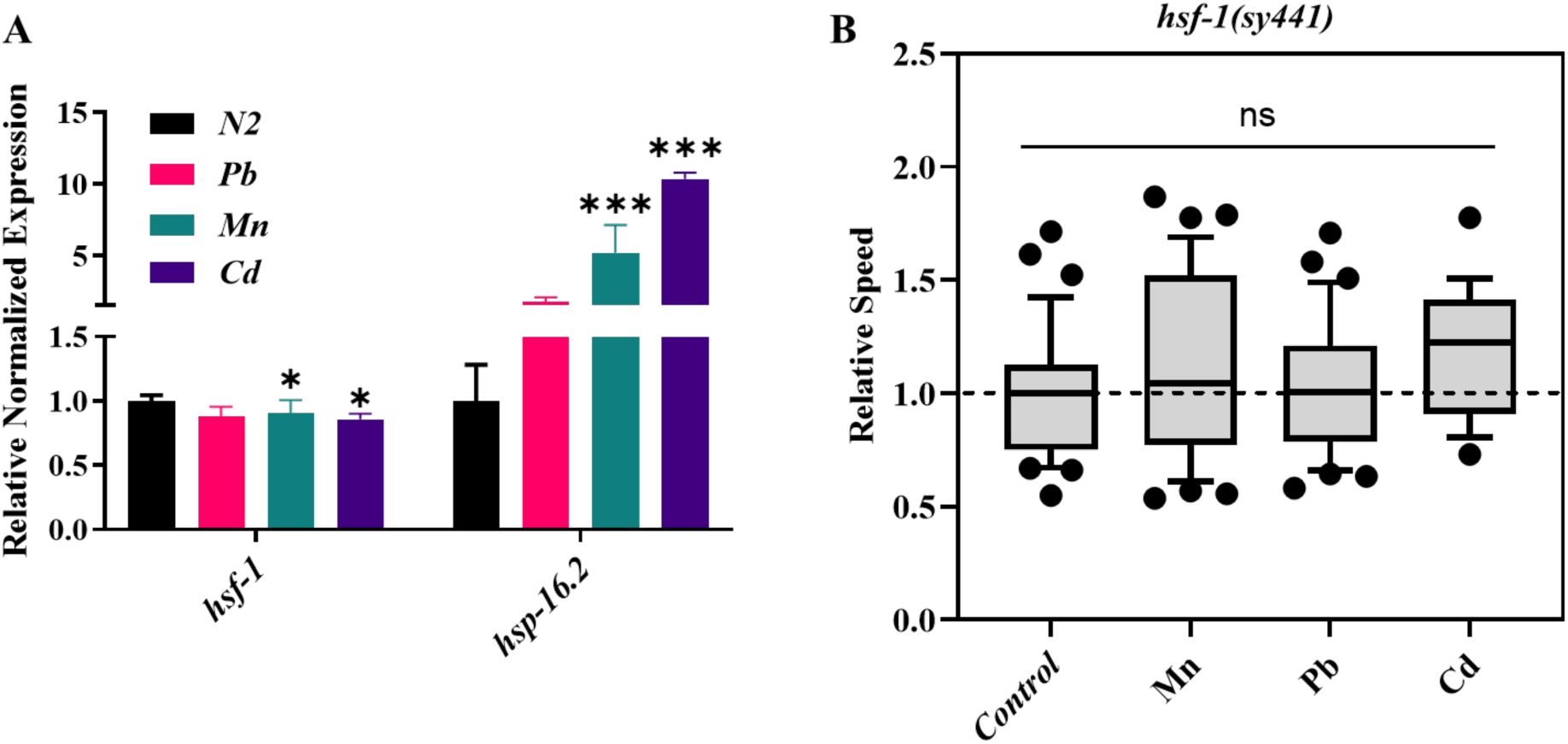
Effect of selected heavy metals of the cytosolic stress response. **A)** RT-qPCR analysis of Mn and Cd exposed worms showing decreased expression of *hsf-1* (p=0.03 and 0.006 respectively) and increased expression of *hsp-16.2* (p<0.001). However, Pb exposure did not significantly alter *hsf-1* (p=0.13494) nor *hsp-16.2* (p=0.578049) expression. Statistical analysis for panel A was done using a one-way ANOVA with Tukey’s post hoc test. **B)** *hsf-1(sy441)* animals were exposed to Pb, Mn, and Cd. No significant difference between any conditions. *p<0.05; **p<0.01; ***p<0.001; ****p<0.0001. **A** is plotted as mean ± SEM

### Post-exposure recovery does not revert electrotaxis defect

Studies have shown that acute exposure to toxic chemicals can have a lasting effect on animals (Benedetto et al., 2010). To examine if exposure paradigm elicits a similar persisting detrimental effect on electrotaxis, Pb and Mn were tested. Both metals caused electrotaxis defects in day-1 adults but in an opposite manner, i.e., Pb induced a hyperactive response characterized by faster speed whereas Mn caused animals to significantly move more slowly compared to control (**Figure 2A**). As an additional follow up to this experiment, we allowed the animals to recover for four days on media plates with no heavy metals but saw no improvement in their movement. Surprisingly, both Pb and Mn exposed worms showed a slower electrotaxis speed (**Figure 6)**. We conclude that exposures of these two metals cause irreversible damage in *C. elegans*.

**Figure 6.**
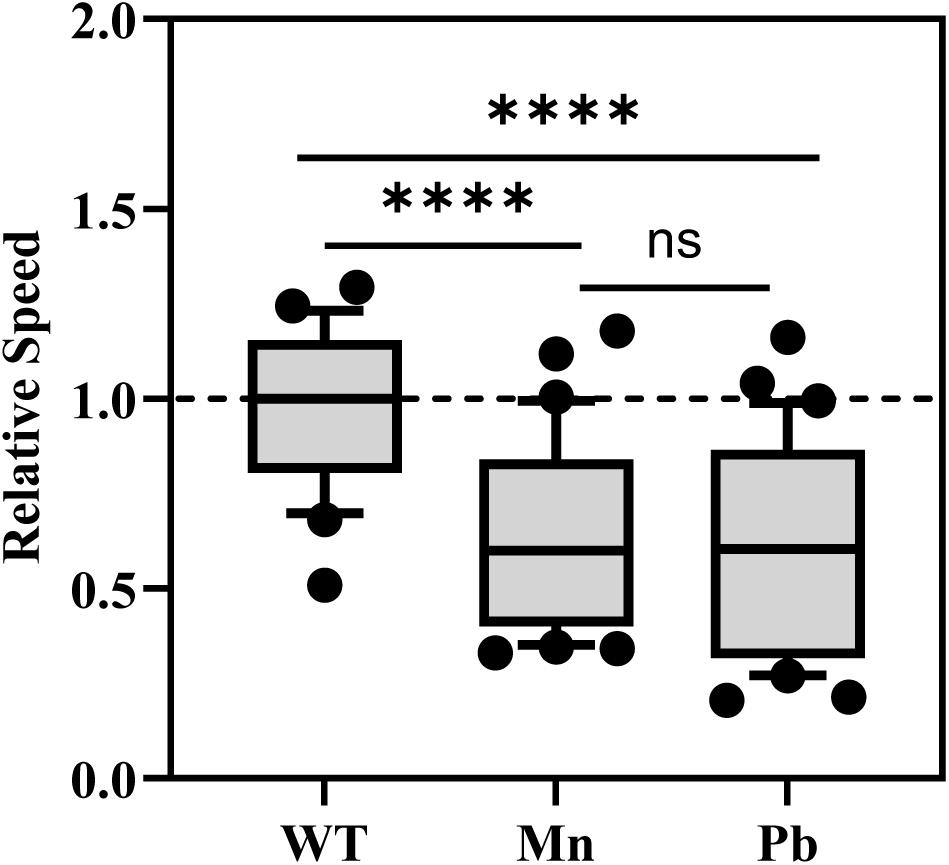
Electrotaxis speed of metal-treated animals following post exposure recovery. Animals were treated with Pb and Mn salts throughout development till day 1 adulthood and allowed to grow on normal, metal-free plates for an additional four days. Following recovery both Mn and Pb significantly reduced the speed of animals. Data pooled from individual replicates (n=20 to 34 animals) per group. Statistical analysis was done using a one-way ANOVA with Dunnett’s post hoc test. *p<0.05; **p<0.01; ***p<0.001; ****p<0.0001.

## DISCUSSION

This study establishes the microfluidic-based electrotaxis assay as a sensitive and quantitative platform for assessing metal-induced neurobehavioral toxicity in *C. elegans*. Unlike traditional locomotion assays that are affected by spontaneous and variable movements that include random pauses, reorientations, omega turns, and backward locomotion (Vidal-Gadea et al., 2012), our approach uses directional electric fields to produce consistent, trackable locomotion and reveal subtle toxicant-induced impairments (Rezai et al., 2010; Tong et al., 2013).

### Electrotaxis defects reflect dose, accumulation, and metal type

We characterized the electrotactic movement of *C. elegans* following exposures to commonly found heavy metals and demonstrate their toxic effects on *C. elegans*’ electrotactic response. Many of the chemicals used in our study such as Hg, MeHg, Fe, Cd, and Ni were reported previously to cause defects in standard plate and liquid assays (Bovio et al., 2021; Colonnello et al., 2020; Fagundez et al., 2015; Ijomone et al., 2020; Klang et al., 2014; Kumar et al., 2015a; Morton and Lamitina, 2013; Moyson et al., 2019; Tang et al., 2019; Wang and Xing, 2008).

We found that the severity of electrotaxis defects correlates with both exposure concentration and metal bioaccumulation. Highly toxic metals such as MeHg, Ag, and Hg impaired electrotaxis at low concentrations and accumulated extensively within the worms. Ag and Hg are known to cause a variety of defects in animals affecting development, behavior, and organ function (Chen et al., 2013; Hunt, 2017). MeHg, in particular, showed ∼33-fold accumulation, consistent with its membrane-permeability and neurotoxicity through LAT1 transport (Helmcke and Aschner, 2010). Cu, though essential, induced behavioral impairment only at high doses. As a physiological mineral, there are endogenous mechanisms in place to regulate Cu homeostasis, including metallochaperones like CUTC-1 (Calafato et al., 2008). Cu’s relatively lower potency in electrotaxis assay may be due to its involvement in normal biological function as an essential component of cytochrome c oxidase, superoxide dismutase, and other conserved enzymes (Halfdanarson et al., 2008). Chronic excesses of Cu in humans, due to regular ingestion or by Cu retention (i.e. Wilson’s disease), are linked to brain and liver damage (Ala et al., 2007). These studies are consistent with our reported results in *C. elegans* here, as well as those by other labs (Vidal-Gadea et al., 2012).

Mn and Pb also impaired electrotaxis, albeit with different signatures. While Mn exposure caused a reduction in speed, Pb-treated animals moved at a faster pace, which may indicate a hyperactive response. While the precise reason of Pb-induced phenotype in worms remains unknown, it is worth mentioning that Pb exposure causes hyperactivity in humans and other systems (Ramirez Ortega et al., 2021; Silbergeld and Goldberg, 1974). The behavioral diversity of these metals may reflect differences in their cellular targets, oxidative stress effects, or neurotransmitter regulation (Akinyemi et al., 2019; Benedetto et al., 2010; Tang et al., 2019). We also examined the post-exposure effect of these metals by allowing worms to recover for four days on clean media. Surprisingly, the animals continued to exhibit abnormal electrotaxis behavior, suggesting irreversible neuronal damage, long-lived stress adaptations, or slow clearance of bioaccumulated metals. The observation that Pb-exposed animals shifted from hyperactivity to reduced speed further underscores the complexity of post-exposure outcomes.

Three of the metals, Ni, Fe, and Cd, did not affect electrotaxis under our chronic exposure conditions. Although these metals are toxic in other contexts, their lack of behavioral effect may reflect insufficient bioavailability or the need for more sensitive movement parameters to detect subtle dysfunctions (Scigajlo, 2015). A longer duration of exposure, i.e., 48 hrs beyond day 1 adulthood, did not cause a change in the speed either. By contrast, Cu caused a slower movement in this assay, revealing differences in sensitivity to metals. Previous studies reported that acute exposure of *C. elegans* to high doses of Ni, Fe, and Cd caused defects in survival, neuronal function and other processes (Colonnello et al., 2020; Hu et al., 2008; Tang et al., 2019; Wang and Xing, 2008). However, these studies used different stages of animals and unique exposure protocols, making direct comparisons to electrotaxis assay difficult (Anderson et al., 2004; Benedetto et al., 2010; Ruszkiewicz et al., 2018). Clearly, more work is needed to compare different metal exposure paradigms in *C. elegans*.

### Neuronal damage parallels behavioral decline, but not always

Behavioral defects were closely mirrored by degeneration of dopaminergic neurons, with metals grouped into three tiers of severity. Group 1 metals (Ag, Hg) caused extensive neuronal damage; Group 2 (Cu, Pb, Mn, Cd) showed intermediate effects; and Group 3 (Ni, Fe) showed little to none. This correlation supports a role for DAergic signaling in electrotaxis regulation (Salam et al., 2013; Taylor et al., 2021), although exceptions such as Cd indicate that additional neuromodulatory systems may be involved or compensate for partial loss. These findings are consistent with other studies showing neurotoxic effects of metals in *C. elegans* (Du and Wang, 2009; Gonzalez-Carter et al., 2017; Hunt, 2017; Ke et al., 2021; Ruszkiewicz et al., 2018).

While the mechanism by which these metals affect neurons is not well understood, one possibility may be increased cellular stress leading to misfolded proteins and protein aggregation. In line with this, Cu was shown to cause an increase in the paralysis of muscle-expressing amyloid beta transgenic worms (Chen et al., 2013). In some cases, metals may directly affect DA signaling, e.g., Pb exposure is reported to interfere with *dat-1* (dopamine transporter) expression resulting in DAergic neuronal dysfunction (Akinyemi et al., 2019). Further investigation into cytosolic stress pathways revealed that while some metals (Mn, Cd) activated *hsp-16.2*, this did not reliably predict behavioral outcomes. Expression of *hsf-1* remained largely unchanged, and *hsf-1* mutant animals showed no enhanced sensitivity to metal-induced defects. This disconnect suggests that the metal-induced electrotaxis phenotype may involve other and/or multiple stress or proteostasis pathways.

### Conclusions

Together, our findings demonstrate that electrotaxis in *C. elegans* is a powerful, rapid, and non-invasive readout of heavy metal toxicity. The assay is capable of detecting both immediate and long-lasting behavioral impairments, thereby offering valuable opportunities for environmental screening, mechanistic toxicology, and high-throughput neurotoxicity testing. Future studies incorporating molecular and electrophysiological analyses could elucidate the distinct pathways through which metals alter neuromuscular function in this genetically tractable animal model.

## MATERIALS AND METHODS

### Strains and culturing

Worms were grown and maintained at 20°C on LB-agar and Modified Youngren’s Only Bactopeptone (MYOB)-agar plates containing *Escherichia coli* OP50 culture using previously described methods (Brenner, 1974). All experiments used age-synchronous populations obtained by bleach treatment. Strains used are N2, *hsf-1(sy441), DY353: bhEx138[pGLC72(Cel-dat-1p::YFP)], MT7988 bas-1(ad446)*.

### Chemicals and treatments

We used the plate-based exposure paradigm which provides efficient bioavailability of chemicals (Vidal-Gadea et al., 2012). All chemicals were obtained from Sigma-Aldrich (St. Louis, MO, USA). All toxicants were first prepared as 200x solutions in 100% DMSO, then diluted 10x with water. 500 μL of the resultant 20x LB agar finally resulting in 1x concentrations of toxicant and 0.5% DMSO.

Nematodes were grown under exposure conditions starting from the L1 larval or egg stage. We chose appropriate concentrations as the highest concentrations of each compound at which chronic exposure would have low to no plate-level phenotypes such as lethality, larval arrest or significant developmental delay. A complete listing of all concentrations tested in this way is shown in **Supplementary Table 1**. Two of the metal salts, AgNO_3_ and HgCl_2_ were tested at three different concentrations (5 μM, 50 μM, and 150 μM). MeHgCl, an organometallic form of Hg, is highly toxic and was used at 2 μM. Cu was tested at a wider range of concentrations (5 μM, 50 μM, 150 μM, and 500 μM), none of which caused an obvious defect at the plate level.

Assays involving five other metals (Cd, Fe, Ni, Pb, and Mn) were initially done at 150 μM concentration. In our plate level analysis, the animals had growth rate, brood size and movement similar to controls with a few exceptions (**Supplementary Table 1**). Cd and Fe caused some delayed growth including arrested L1 larvae. Higher concentrations of both of these chemicals and Ni were detrimental as L1 larvae showed growth arrest. The movement tracks and brood size of the worms on day 1 adulthood were also very similar to untreated controls.

Two of the metals, Fe and Cd, were also tested at other concentrations. Animals exposed to 300 μM of FeSO4 showed obvious defects on plates (i.e., growth arrest), which precluded any further analysis. Cd was less toxic at 100 μM but still caused severe growth arrest. However, animals exposed to 50 μM Cd showed a mild growth defect at the plate level but otherwise appeared normal during adulthood, thereby allowing us to test their electrotaxis responses.

We first observed *hsf-1* animals at a plate level and found that those exposed to Cd had arrested at L1 larval stage, but some were still able to reach adulthood. In the case of Pb exposure, animals were developmentally delayed by a few hours and Mn did not influence the animals overall. All together the animals looked visually healthy and had normal movement tracks on the plate.

### Electrotaxis assay

Microfluidic channels were fabricated as previously described (Rezai et al., 2010). Adult worms were washed off their culture plates, cleaned, and suspended in M9 buffer. Animals were then aspirated into the channel using the syringe pump. Individual animals were isolated by adjusting the tubes’ relative height to hydrostatically manipulate the flow of M9 through the channel. Both tubes were then laid flat at the same elevation to eliminate pressure-induced flow. Next, a 3 V/cm DC electric field was applied and the animal’s resultant behavior recorded by camera. Locomotory data was later extracted from recorded videos either manually using NIH ImageJ (http://rsbweb.nih.gov/ij/) or automatically with custom MATLAB-based tracking software (Scigajlo, 2015; Tong et al., 2013). Toxicant-exposed animals were grown for 69 h at 20°C before scoring. At least three batches of 10 animals each were scored for each condition. Speed data was normalized to the median of the control.

### Brood size, growth and lifespan assays

Reproductive capacity of toxicant-exposed animals was determined by counting the hatched progeny of isolated hermaphrodites. Five L4-stage larvae were picked onto fresh spiked MYOB plates and incubated for an additional 96 h at 20°C for maturation and egg-laying; afterwards, the number of offspring was estimated by suspending the population in M9 and counting the animals in aliquots of the suspension. In the cases of 50 and 150 μM Ag and Hg, nematodes were grown for 2 weeks instead of 6 days to account for these animals’ slower growth, and progeny were counted and removed from plates daily. The resultant progeny on two batches of four plates per test condition were quantified in this way.

Body length of metal-exposed nematodes was determined after 69 h of growth at 20°C on toxicant-spiked MYOB plates containing food. Measurements were made from photographs of animals anesthetized through placement in a 15-μL drop of 30 mM NaN_3_ in M9, which lay on a solidified pad of 4% agar on a glass slide. Photographs were taken and analyzed with a Hamamatsu ORCA-AG camera on a Nikon Eclipse 80i Nomarski microscope, using NIS-Elements BR software version 3.0 (www.nis-elements.com). At least two batches of 10 animals each were measured for each test condition.

For lifespan assays two batches of 20 age synchronous L1 larvae per condition were transferred into four fresh spiked MYOB plates (five animals per plate) and grown at 20°C. Viability of animals was checked each day under a stereomicroscope. If immobile, animals were checked for viability by tapping the plate and gently touching their bodies with a platinum pick. Nematodes were transferred to fresh spiked MYOB plates after 120 h, 168 h, and 216 h to prevent contamination of parent animals by offspring.

### Determination of metal content

Approximately 10,000 age-synchronous L1 larvae per condition were transferred into fresh spiked MYOB plates and grown at 20°C for 69 h, at the conclusion of which both live and dead animals were collected and washed twice with deionized water. Two biological replicates were prepared for each exposure. Untreated worms were used as controls. The pelleted pool of worms was frozen at −80°C for further processing at Actlabs (Ancaster, ON, Canada). Each sample was weighed into a 50-mL centrifuge tube followed by the addition of 2 mL of concentrated HNO_3_. Centrifuge tubes were placed in a boiling water bath for 1 h to digest samples. The fully digested content in each tube was then diluted to 50 mL with water and analyzed by inductively coupled plasma-mass spectrometry (ICP-MS) using QOP Hydrogeo Rev. 6.6. In addition to testing worms, we also measured metal concentrations in the exposure media.

The elemental contents of Ag, Hg, and MeHg were measured as a test case to determine the effectiveness of our agar plate-based exposure paradigm. The results revealed that while Hg was recovered with somewhat lower efficiency (71% for HgCl_2_, and 37% for MeHgCl), possibly due to speciation and precipitation, Ag was fully recoverable.

### Neuronal phenotype analyses

Animals were mounted on 2% agar pad containing glass slides. Before placing the cover slip, they were anesthetized using 30 mM NaN_3_. GFP fluorescence was visualized using a Zeiss Observer Z1 microscope equipped with an Apotome 2 and X-Cite R 120LED fluorescence illuminator. The *dat-1p::YFP* strain was used to visualize CEP and ADE dopaminergic neurons (Taylor et al., 2021). The percentage of animals with defects was plotted.

### Basal Slowing Response Assay

The protocol was adapted from a published study (Sawin et al., 2000). Body Bend was measured for 20 secs 3 times per worm. 200 μL of bacteria was plated to seed half of the plate. Worms were transferred to the area without food and then allowed to crawl to the side of the plate with food. Day 3 adult animals were used. Experiment was repeated in two independent trials(replicates) using 8 animals per trial for each metal.

### RNA extraction and RT-qPCR

In brief, the RNA extraction protocol was adapted from a published study (Taylor et al., 2024). Animals were bleach synchronized and then harvested at day 1 adult stage. RNA was extracted using Trizol/ chloroform method and then samples were DNAse treated with TURBO DNA-free™ Kit (Catalog Number:AM1907, ThermoFisher Scientific), and cDNA synthesis was made using the SensiFAST cDNA Synthesis Kit (Catalog Number BIO-65053, Meridian Bioscience). RT-qPCR was performed in triplicate using the Bio-Rad cycler CFX 96 and the SensiFAST SYBR Green Kit (Catalog Number BIO-98005, BIOLINE, USA). Expression levels were normalized to the housekeeping gene *pmp-3.* RT-qPCR graphs were plotted as mean ± SEM.

### Post-exposure recovery

Animals were grown in the presence of metal salts from egg stage till day 1 adulthood. The animals were washed thoroughly and move to normal OP50 culture plates. They were transferred every other day until day 5 adulthood.

### Data analysis

Graphpad Prism 9 was used to perform statistical analysis. Electrotaxis speed data was plotted in box plots and compared with the non-parametric Mann-Whitney test. Lifespan data was assessed with the Kaplan-Meier method and compared with the log-rank test. All other data was analyzed with Student’s *t*-test. For all assays, data from all repeats (minimum 3) were pooled and analyzed together.

## ACKNOWLEDGEMENTS

We thank Anne Xia and Dr. Pouya Rezai for discussions and assistance with some of the experiments. Some strains were provided by the CGC, which is funded by NIH Office of Research Infrastructure Programs. We are grateful to Drs. Chris Wood and James McNulty for providing some of the chemicals and advice. This work was supported by grants from the Ontario Ministry of Research and Innovation (BPG), Collaborative Health Research Project, co-funded by the Natural Sciences and Engineering Research Council of Canada and the Canadian Institutes of Health Research (PRS, BPG and RKM), and Natural Sciences and Engineering Research Council of Canada Discovery grant (BPG).

## List of Tables

**Table 1.**
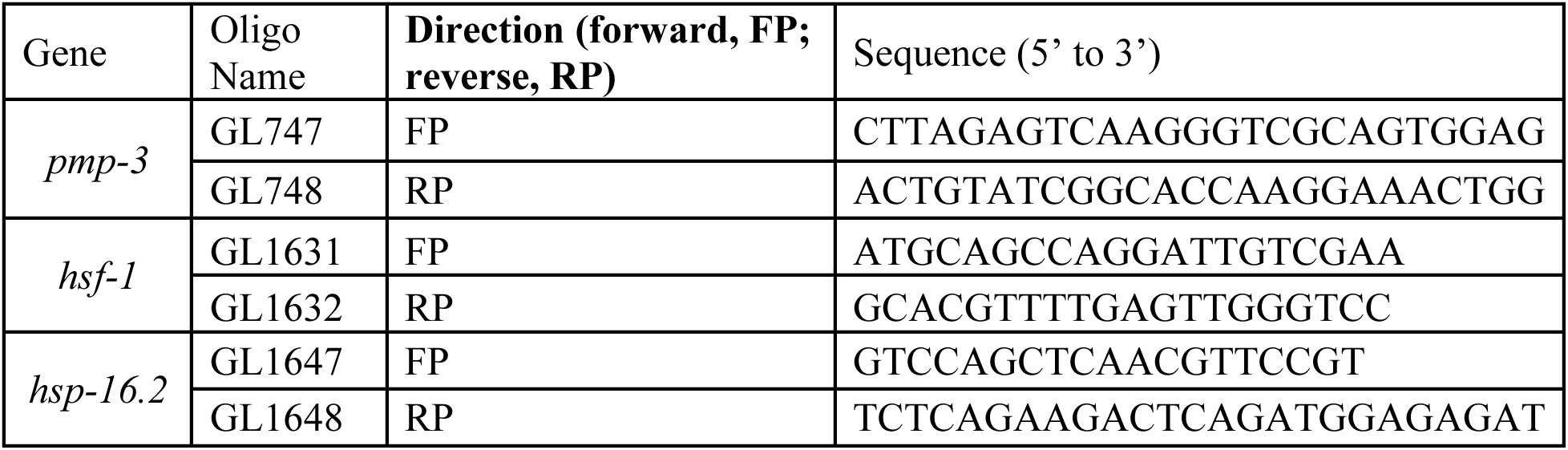
List of primers used in RT-qPCR analysis.

## SUPPLMENTARY MATERIALS

**Supplementary Figure 1.**
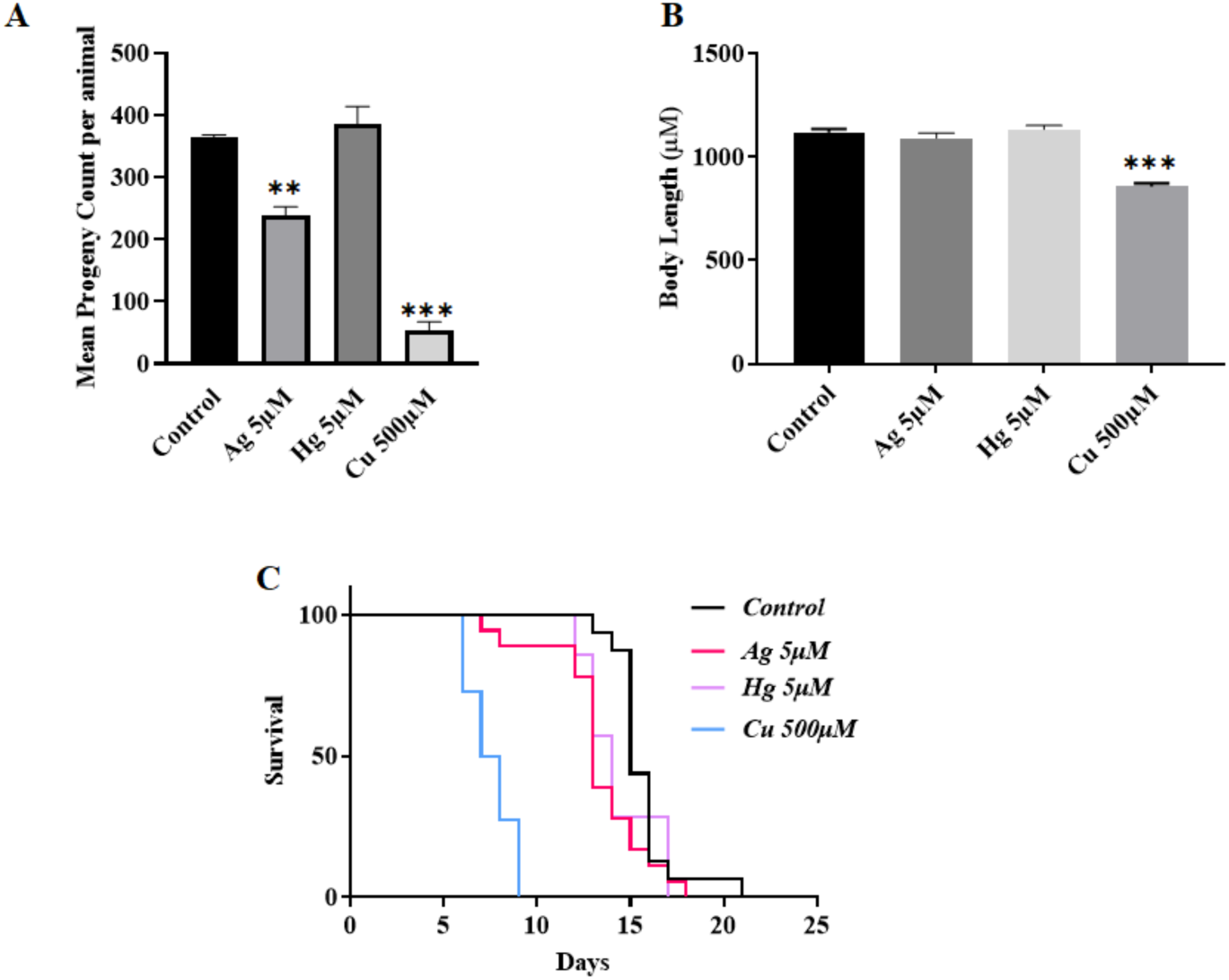
Influence of chronic Ag, Hg, and Cu exposures on reproduction, body length and lifespan. **A)** reproductive capacity of *C. elegans*. At 5 μM Ag caused 36% reduction in brood size, however 5 μM Hg had no effect. Increasing the concentration by 10 fold, i.e. 50 μM, both Ag and Hg caused 99% and 87% reduced brood size, respectively. **B)** Body length was also reduced following exposure to Ag and Hg. Ag 50 μM: 39% length reduction, Ag 150 μM: 56% reduction, Hg 50 μM: 62% reduction, Hg 150 μM: 77% reduction. **C)** Influence of chronic metal exposure on lifespan. 5 μM Ag and Hg had no obvious effect on the lifespan. Cu 500 μM significantly reduced the lifespan of animals.Data is pooled from independent replicates (n > 30 animals). Statistical analysis for panel A and B was done using a one-way ANOVA with Dunnett’s post hoc test. The lifespan data for C were plotted as survival curves which were estimated using the Kaplan-Meier test followed by the log-rank test for group differences. *p<0.05; **p<0.01; ***p<0.001; ****p<0.0001. **A** and **B** are plotted as mean ± SEM

**Supplementary Figure 2.**
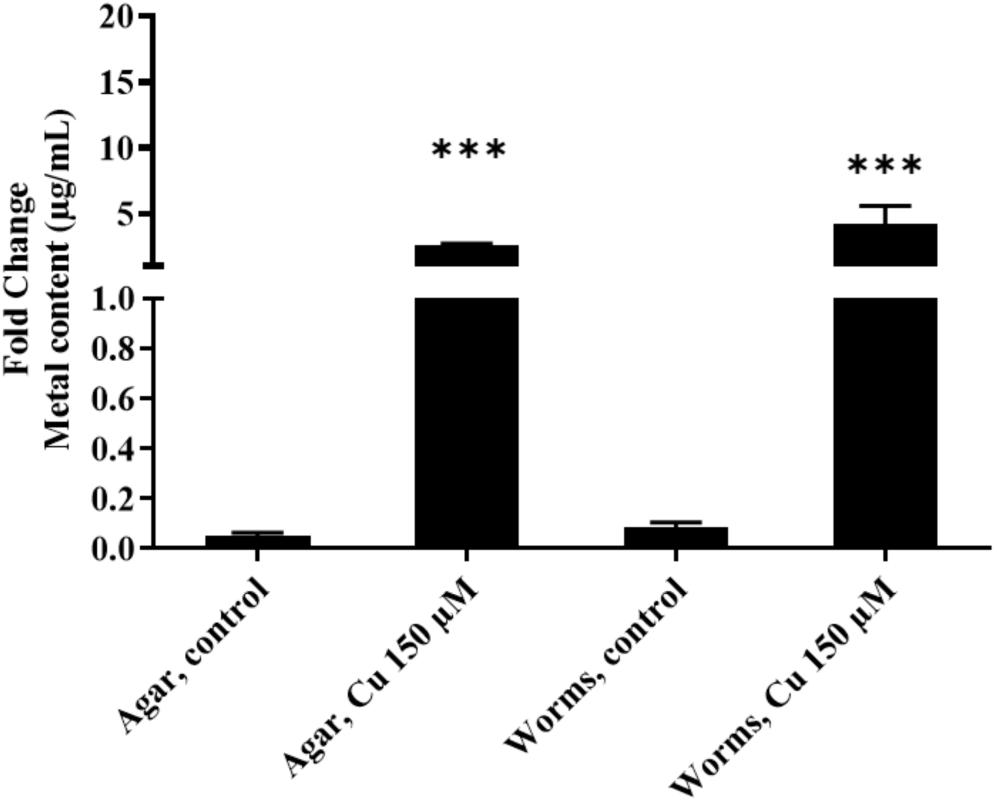
Metal content in *C. elegans* following 69 hrs Cu exposure. Elemental content was measured as a function of total sample volume. Metal content in worms is significantly higher than in control following treatment with Cu 150 μM. Student’s unpaired t-test was used. *p<0.05; **p<0.01; ***p<0.001; ****p<0.0001. Plotted as mean ± SEM

**Supplementary Figure 3.**
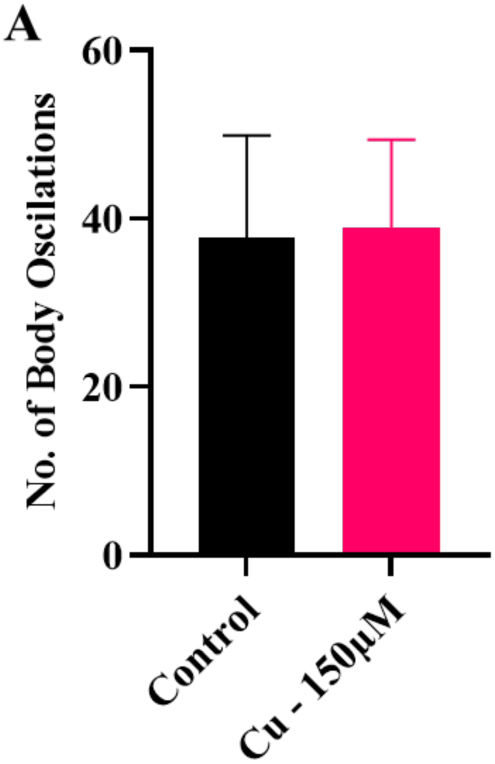
Influence of chronic Cu exposure on body oscillation. The data was produced by the MATLab software. Cu at 150µM caused no significant changes in body oscillations of animals. Plotted as mean ± SD

**Supplementary Table 1.** Plate-level observations of animals exposed to different concentrations of metal salts. The phenotypes included slower developmental growth, arrest during the L1 larval stage (including lethality), and abnormal appearance (slower movement or no movement, reduced foraging). Animals were scored as normal (no obvious difference from untreated controls), intermediate (phenotype observed in some animals), and defective (phenotype clearly visible in most or all animals). The dash (-) indicates animals that couldn’t be analyze due to growth arrest.

|  | **L1 arrest** | **Developmental growth** | **Abnormal appearance** |
| --- | --- | --- | --- |
| **Ni 150μM** | Intermediate | Normal | Normal |
| **Pb - 150μM** | Normal | Normal | Normal |
| **Mn 150μM** | Normal | Normal | Normal |
| **Fe 150μM** | Intermediate | Normal | Normal |
| **Fe 300μM** | Defective | - | Intermediate |
| **Cd 50μM** | Intermediate | Intermediate | Normal |
| **Cd 100μM** | defective | defective | intermediate |
| **Cd 150μM** | defective | - | - |
| **Cu 150μM** | Normal | Normal | Normal |
| **Cu 500μM** | Normal | Normal | Normal |

## REFERENCES

Akinyemi, A.J., Miah, M.R., Ijomone, O.M., Tsatsakis, A., Soares, F.A.A., Tinkov, A.A., Skalny, A.V., Venkataramani, V., Aschner, M., 2019. Lead (Pb) exposure induces dopaminergic neurotoxicity in Caenorhabditis elegans: Involvement of the dopamine transporter. Toxicol Rep 6, 833–840.

Ala, A., Walker, A.P., Ashkan, K., Dooley, J.S., Schilsky, M.L., 2007. Wilson’s disease. Lancet 369, 397–408.

Anderson, G.L., Cole, R.D., Williams, P.L., 2004. Assessing behavioral toxicity with Caenorhabditis elegans. Environ Toxicol Chem 23, 1235–1240.

Asharani, P.V., Hande, M.P., Valiyaveettil, S., 2009. Anti-proliferative activity of silver nanoparticles. BMC Cell Biol 10, 65.

Bar-Ilan, O., Albrecht, R.M., Fako, V.E., Furgeson, D.Y., 2009. Toxicity assessments of multisized gold and silver nanoparticles in zebrafish embryos. Small 5, 1897–1910.

Benedetto, A., Au, C., Avila, D.S., Milatovic, D., Aschner, M., 2010. Extracellular dopamine potentiates mn-induced oxidative stress, lifespan reduction, and dopaminergic neurodegeneration in a BLI-3-dependent manner in Caenorhabditis elegans. PLoS Genet 6.

Bovio, F., Sciandrone, B., Urani, C., Fusi, P., Forcella, M., Regonesi, M.E., 2021. Superoxide dismutase 1 (SOD1) and cadmium: A three models approach to the comprehension of its neurotoxic effects. Neurotoxicology 84, 125–135.

Brenner, S., 1974. The genetics of *Caenorhabditis elegans*. Genetics 77, 71–94.

Calafato, S., Swain, S., Hughes, S., Kille, P., Sturzenbaum, S.R., 2008. Knock down of Caenorhabditis elegans cutc-1 exacerbates the sensitivity toward high levels of copper. Toxicol Sci 106, 384–391.

Chen, P., Martinez-Finley, E.J., Bornhorst, J., Chakraborty, S., Aschner, M., 2013. Metal-induced neurodegeneration in C. elegans. Front Aging Neurosci 5, 18.

Clarkson, T.W., Magos, L., 2006. The toxicology of mercury and its chemical compounds. Crit Rev Toxicol 36, 609–662.

Colonnello, A., Aguilera-Portillo, G., Rubio-Lopez, L.C., Robles-Banuelos, B., Rangel-Lopez, E., Cortez-Nunez, S., Evaristo-Priego, Y., Silva-Palacios, A., Galvan-Arzate, S., Garcia-Contreras, R., Tunez, I., Chen, P., Aschner, M., Santamaria, A., 2020. Correction to: Comparing the Neuroprotective Effects of Caffeic Acid in Rat Cortical Slices and Caenorhabditis elegans: Involvement of Nrf2 and SKN-1 Signaling Pathways. Neurotox Res 37, 779.

Drake, P.L., Hazelwood, K.J., 2005. Exposure-related health effects of silver and silver compounds: a review. Ann Occup Hyg 49, 575–585.

Du, M., Wang, D., 2009. The neurotoxic effects of heavy metal exposure on GABAergic nervous system in nematode Caenorhabditis elegans. Environ Toxicol Pharmacol 27, 314–320.

Fagundez, D.d.A., Câmara, D.F., Salgueiro, W.G., Noremberg, S., Luiz Puntel, R., Piccoli, J.E., Garcia, S.C., da Rocha, J.B.T., Aschner, M., Ávila, D.S., 2015. Behavioral and dopaminergic damage induced by acute iron toxicity in Caenorhabditis elegans. Toxicology Research 4, 878–884.

Forsyth, D.S., Casey, V., Dabeka, R.W., McKenzie, A., 2004. Methylmercury levels in predatory fish species marketed in Canada. Food Addit Contam 21, 849–856.

Genchi, G., Carocci, A., Lauria, G., Sinicropi, M.S., Catalano, A., 2020. Nickel: Human Health and Environmental Toxicology. Int J Environ Res Public Health 17.

Gonzalez-Carter, D.A., Leo, B.F., Ruenraroengsak, P., Chen, S., Goode, A.E., Theodorou, I.G., Chung, K.F., Carzaniga, R., Shaffer, M.S., Dexter, D.T., Ryan, M.P., Porter, A.E., 2017. Silver nanoparticles reduce brain inflammation and related neurotoxicity through induction of H(2)S-synthesizing enzymes. Sci Rep 7, 42871.

Hadrup, N., Sharma, A.K., Loeschner, K., 2018. Toxicity of silver ions, metallic silver, and silver nanoparticle materials after in vivo dermal and mucosal surface exposure: A review. Regul Toxicol Pharmacol 98, 257–267.

Halfdanarson, T.R., Kumar, N., Li, C.Y., Phyliky, R.L., Hogan, W.J., 2008. Hematological manifestations of copper deficiency: a retrospective review. Eur J Haematol 80, 523–531.

Hart, A.C., 2006. Behavior. Wormbook ed. The C. elegans Research Community, WormBook, 10.1895/wormbook.1.87.1, http://www.wormbook.org.

Helmcke, K.J., Aschner, M., 2010. Hormetic effect of methylmercury on Caenorhabditis elegans. Toxicol Appl Pharmacol 248, 156–164.

Hong, Y.S., Kim, Y.M., Lee, K.E., 2012. Methylmercury exposure and health effects. J Prev Med Public Health 45, 353–363.

Hu, Y.O., Wang, Y., Ye, B.P., Wang, D.Y., 2008. Phenotypic and behavioral defects induced by iron exposure can be transferred to progeny in Caenorhabditis elegans. Biomed Environ Sci 21, 467–473.

Hunt, P.R., 2017. The C. elegans model in toxicity testing. J Appl Toxicol 37, 50–59.

Ijomone, O.M., Miah, M.R., Akingbade, G.T., Bucinca, H., Aschner, M., 2020. Nickel-Induced Developmental Neurotoxicity in C. elegans Includes Cholinergic, Dopaminergic and GABAergic Degeneration, Altered Behaviour, and Increased SKN-1 Activity. Neurotox Res 37, 1018–1028.

Jaishankar, M., Tseten, T., Anbalagan, N., Mathew, B.B., Beeregowda, K.N., 2014. Toxicity, mechanism and health effects of some heavy metals. Interdiscip Toxicol 7, 60–72.

Ju, J., Ruan, Q., Li, X., Liu, R., Li, Y., Pu, Y., Yin, L., Wang, D., 2013. Neurotoxicological evaluation of microcystin-LR exposure at environmental relevant concentrations on nematode Caenorhabditis elegans. Environ Sci Pollut Res Int 20, 1823–1830.

Ke, T., Prince, L.M., Bowman, A.B., Aschner, M., 2021. Latent alterations in swimming behavior by developmental methylmercury exposure are modulated by the homolog of tyrosine hydroxylase in Caenorhabditis elegans. Neurotoxicol Teratol 85, 106963.

Kim, J., Song, H., Heo, H.R., Kim, J.W., Kim, H.R., Hong, Y., Yang, S.R., Han, S.S., Lee, S.J., Kim, W.J., Hong, S.H., 2017. Cadmium-induced ER stress and inflammation are mediated through C/EBP-DDIT3 signaling in human bronchial epithelial cells. Exp Mol Med 49, e372.

Klang, I.M., Schilling, B., Sorensen, D.J., Sahu, A.K., Kapahi, P., Andersen, J.K., Swoboda, P., Killilea, D.W., Gibson, B.W., Lithgow, G.J., 2014. Iron promotes protein insolubility and aging in C. elegans. Aging (Albany NY) 6, 975–991.

Kumar, R., Pradhan, A., Khan, F.A., Lindstrom, P., Ragnvaldsson, D., Ivarsson, P., Olsson, P.E., Jass, J., 2015a. Comparative Analysis of Stress Induced Gene Expression in Caenorhabditis elegans following Exposure to Environmental and Lab Reconstituted Complex Metal Mixture. PloS one 10, e0132896.

Kumar, V., Kalita, J., Misra, U.K., Bora, H.K., 2015b. A study of dose response and organ susceptibility of copper toxicity in a rat model. J Trace Elem Med Biol 29, 269–274.

Liu, X.D., Thiele, D.J., 1996. Oxidative stress induced heat shock factor phosphorylation and HSF-dependent activation of yeast metallothionein gene transcription. Genes Dev 10, 592–603.

Martinez-Finley, E.J., Aschner, M., 2011. Revelations from the Nematode Caenorhabditis elegans on the Complex Interplay of Metal Toxicological Mechanisms. Journal of toxicology 2011, 895236.

McElwee, M.K., Freedman, J.H., 2011. Comparative toxicology of mercurials in Caenorhabditis elegans. Environ Toxicol Chem 30, 2135–2141.

Mo, Y., Jiang, M., Zhang, Y., Wan, R., Li, J., Zhong, C.J., Li, H., Tang, S., Zhang, Q., 2019. Comparative mouse lung injury by nickel nanoparticles with differential surface modification. J Nanobiotechnology 17, 2.

Morton, E.A., Lamitina, T., 2013. Caenorhabditis elegans HSF-1 is an essential nuclear protein that forms stress granule-like structures following heat shock. Aging Cell 12, 112–120.

Moyson, S., Town, R.M., Vissenberg, K., Blust, R., 2019. The effect of metal mixture composition on toxicity to C. elegans at individual and population levels. PloS one 14, e0218929.

Ndayisaba, A., Kaindlstorfer, C., Wenning, G.K., 2019. Iron in Neurodegeneration - Cause or Consequence? Front Neurosci 13, 180.

Neal, A.P., Guilarte, T.R., 2010. Molecular neurobiology of lead (Pb(2+)): effects on synaptic function. Mol Neurobiol 42, 151–160.

Rahman, M.F., Wang, J., Patterson, T.A., Saini, U.T., Robinson, B.L., Newport, G.D., Murdock, R.C., Schlager, J.J., Hussain, S.M., Ali, S.F., 2009. Expression of genes related to oxidative stress in the mouse brain after exposure to silver-25 nanoparticles. Toxicol Lett 187, 15–21.

Raj, V., Nair, A., Thekkuveettil, A., 2021. Manganese exposure during early larval stages of C. elegans causes learning disability in the adult stage. Biochem Biophys Res Commun 568, 89–94.

Ramirez Ortega, D., Gonzalez Esquivel, D.F., Blanco Ayala, T., Pineda, B., Gomez Manzo, S., Marcial Quino, J., Carrillo Mora, P., Perez de la Cruz, V., 2021. Cognitive Impairment Induced by Lead Exposure during Lifespan: Mechanisms of Lead Neurotoxicity. Toxics 9.

Rezai, P., Siddiqui, A., Selvaganapathy, P.R., Gupta, B.P., 2010. Electrotaxis of Caenorhabditis elegans in a microfluidic environment. Lab Chip 10, 220–226.

Roh, J.Y., Sim, S.J., Yi, J., Park, K., Chung, K.H., Ryu, D.Y., Choi, J., 2009. Ecotoxicity of silver nanoparticles on the soil nematode Caenorhabditis elegans using functional ecotoxicogenomics. Environ Sci Technol 43, 3933–3940.

Ruszkiewicz, J.A., Pinkas, A., Miah, M.R., Weitz, R.L., Lawes, M.J.A., Akinyemi, A.J., Ijomone, O.M., Aschner, M., 2018. C. elegans as a model in developmental neurotoxicology. Toxicol Appl Pharmacol 354, 126–135.

Salam, S., Ansari, A., Amon, S., Rezai, P., Selvaganapathy, P.R., Mishra, R.K., Gupta, B.P., 2013. A microfluidic phenotype analysis system reveals function of sensory and dopaminergic neuron signaling in C. elegans electrotactic swimming behavior. Worm 2, e24558.

Sawin, E.R., Ranganathan, R., Horvitz, H.R., 2000. C. elegans locomotory rate is modulated by the environment through a dopaminergic pathway and by experience through a serotonergic pathway. Neuron 26, 619–631.

Scigajlo, A., 2015. Automated Nematode Tracking System, Electrical & Computer Engineering. McMaster University, Hamilton, p. 81.

Shen, L., Xiao, J., Ye, H., Wang, D., 2009. Toxicity evaluation in nematode Caenorhabditis elegans after chronic metal exposure. Environ Toxicol Pharmacol 28, 125–132.

Silbergeld, E.K., Goldberg, A.M., 1974. Hyperactivity: a lead-induced behavior disorder. Environ Health Perspect 7, 227–232.

Tang, B., Tong, P., Xue, K.S., Williams, P.L., Wang, J.S., Tang, L., 2019. High-throughput assessment of toxic effects of metal mixtures of cadmium(Cd), lead(Pb), and manganese(Mn) in nematode Caenorhabditis elegans. Chemosphere 234, 232–241.

Taylor, S.K.B., Hartman, J.H., Gupta, B.P., 2024. The neurotrophic factor MANF regulates autophagy and lysosome function to promote proteostasis in Caenorhabditis elegans. Proc Natl Acad Sci U S A 121, e2403906121.

Taylor, S.K.B., Minhas, M.H., Tong, J., Selvaganapathy, P.R., Mishra, R.K., Gupta, B.P., 2021. C. elegans electrotaxis behavior is modulated by heat shock response and unfolded protein response signaling pathways. Sci Rep 11, 3115.

Tong, J., Rezai, P., Salam, S., Selvaganapathy, P.R., Gupta, B.P., 2013. Microfluidic-based electrotaxis for on-demand quantitative analysis of Caenorhabditis elegans’ locomotion. Journal of visualized experiments : JoVE, e50226.

Uenishi, R., Gong, P., Suzuki, K., Koizumi, S., 2006. Cross talk of heat shock and heavy metal regulatory pathways. Biochem Biophys Res Commun 341, 1072–1077.

Vidal-Gadea, A.G., Davis, S., Becker, L., Pierce-Shimomura, J.T., 2012. Coordination of behavioral hierarchies during environmental transitions in Caenorhabditis elegans. Worm 1, 5–11.

Wang, D., Xing, X., 2008. Assessment of locomotion behavioral defects induced by acute toxicity from heavy metal exposure in nematode Caenorhabditis elegans. J Environ Sci (China) 20, 1132–1137.

Wang, S., You, M., Wang, C., Zhang, Y., Fan, C., Yan, S., 2020. Heat shock pretreatment induced cadmium resistance in the nematode Caenorhabditis elegans is depend on transcription factors DAF-16 and HSF-1. Environ Pollut 261, 114081.

Wazir, S.M., Ghobrial, I., 2017. Copper deficiency, a new triad: anemia, leucopenia, and myeloneuropathy. J Community Hosp Intern Med Perspect 7, 265–268.

Wu, Q., Qu, Y., Li, X., Wang, D., 2012. Chromium exhibits adverse effects at environmental relevant concentrations in chronic toxicity assay system of nematode Caenorhabditis elegans. Chemosphere 87, 1281–1287.

